# Tumor Origin Detection with Tissue-Specific miRNA and DNA methylation Markers

**DOI:** 10.1101/090746

**Authors:** Wei Tang, Shixiang Wan, Quan Zou

## Abstract

**Motivation:** Cancer of unknown primary origin constitutes 3-5% of all human malignancies. Patients with these carcinomas present with metastases without an established primary site, which may not be found even by thorough histological search methods. Patients with cancer of unknown primary origin always have poor prognosis and hardly have efficient treatment since most cancers respond well to specific chemotherapy or hormone drugs. Many studies have proposed classifiers based on miRNAs or mRNAs to predict the tumor origins, but few study focus on high-dimensional DNA methylation profiles.

**Results:** We introduced three classifiers with novel feature selection algorithm combined with random forest to effectively identify highly tissue-specific epigenetics biomarkers such as microRNAs and CpG sites, which can help us predict the origin site of tumors. This algorithm, incorporating differential analysis and descending dimension algorithm, was applied on 14 histological tissues and over 5000 samples based on miRNA expression and DNA methylation profiles to assign given primary tumor to its origin tissue. Our study shows all of these three classifiers have an overall accuracy of 87.78% (72.55%-97.54%) based on miRNA datasets and an accuracy of 96.43% (MRMD: 87.85%-99.76%) or 97.06% (PCA: 92.44%-100%) based on DNA methylation datasets on predicting the origin of tumors and suggests that the biomarkers we selected can efficiently predict the origin of tumors and allow the clinicians to avoid adjuvant systemic therapy or to choose less aggressive therapeutic options. We also developed a user-friendly webserver which enables users to predict the origin site of tumors by uploading the miRNAs expression or DNA methylation profiles of those cancers.

**Availability:** The webserver, data, and code are accessible free of charge at http://server.malab.cn/MMCOP/

**Supplementary information:** Supplementary data are available at *Bioinformatics* online.

## 1 Introduction

In a general medical oncology service, Metastatic Cancer of Unknown Primary Origin (CUP) whose origin site of a tumor cannot be readily identified, which may account about for 3%-5% in the new cancer cases (Chaffer and Weinberg, 2011; Greco and Hainsworth, 2006; Gupta and Massagué, 2006; Hodi, et al., 2010; Joyce and Pollard, 2009). The origin site of CUP can also remain equivocal, even after thorough complete physical examination including pelvic and rectal examination (ADLER, et al., 1999; Willett, et al., 2009), full blood count and biochemistry, urinalysis and stool occult blood testing, histological evaluation of biopsy material with the use of immunohistochemistry (Kwak, et al., 2010), radiological examination (Suzuki, et al., 2011), computed tomography (CT) of the abdomen (Giacchetti, et al., 2000; Investigators, 2006) and, in certain cases, mammography (Chlebowski, et al., 2010) fail to identify the primary site. This disease manifestation with highly heterogeneous characteristic is also one of the 10 most frequent cancers worldwide (Chaffer and Weinberg, 2011). CUP also represents a clinically diverse group, typically presenting with moderately to well-differentiated adenocarcinoma, or undifferentiated to poorly differentiated tumors, involving multiple organs such as liver, bone, lung, lymph nodes and breast (Greco and Hainsworth, 2006). Patients who are diagnosed as CUP with elusive origin site often represent an incommensurate proportion of cancer deaths or a very poor median survival, often measured in months, the average survival time of these patients would be only about 9 to 12 months with a very low survival rate after diagnosis (Daugaard, et al., 2009). Even though the survival rate depends on various factors such as the cancer cell type, where the cancer is found, how far the cancer has spread, the treatments received, and how well the cancer responds to treatment, the main reason why there exist a high mortality among CUP patients can be account for by the misclassification of unknown tumor origin. A large proportion of cases remain undiagnosed or mistakenly diagnosed, with the result that therapy cannot be matched to their specific disease, particularly for certain cancers that respond well to specific chemotherapy or hormone drugs.

The rapidly development of next-generation sequencing (NGS) technology (Hanahan and Weinberg, 2011; McKenna, et al., 2010) has not only greatly advanced the generation of vast high-throughput sequence data and revolutionized the human genomics research through the analysis on linking the phenotype to its genomics, including DNA microarray (ChIP-chip) and RNA sequencing (RNA-seq) (Hormozdiari, et al., 2009; Kozubek, et al., 2013), but also enhances epigenetic studies with high coverage density and flexibility (Hurd and Nelson, 2009; Ku, et al., 2011). Particularly, a major extension of the previous Infinium HumanMethylation27 BeadChip, called Infinium HumanMethylation450 (Infinium Methylation 450K; Illumina, Inc. CA, USA) was developed as a powerful technique in terms of agentia costs, sample throughput and coverage (Aryee, et al., 2014; Bibikova, et al., 2009; Dedeurwaerder, et al., 2011; Sandoval, et al., 2011). It holds a great promise for a better understanding of the epigenetic component in health and disease and promotes a more comprehensive view of methylation patterns at single-base resolution across the genome with the help of whole-genome bisulfite sequencing (WGBS) (Xi and Li, 2009) which leverages the power of NGS (Chen, et al., 2013; Kim, et al., 2011; Lister, et al., 2009). Since developing therapeutic strategies, especially those involving specific therapy, would be much more challenging in CUP cases, and use of sequencing microarrays can promise to ultimately be of help in this regard. Tumor classifications based on gene expression designed for clinical application to CUP, particularly on miRNAs expression profiles, which are small non-protein-coding RNAs (Bartel, 2009) that have been much found to regulate the expression of gene involved in many biological processes, such as cell proliferation, cell death and differentiation (Bartel, 2009; Hayashita, et al., 2005; Hwang and Mendell, 2006), had been proposed to predict the origin of tumors (Budhu, et al., 2008; Heinzelmann, et al., 2011; Rosenfeld, et al., 2008). For example, Rolf et al (Søkilde, et al., 2014) developed a miRNA-based classifier which mainly involved feature selection embedded in the Least Absolute Shrinkage and Selection Operator (LASSO) classification algorithm, this classifier has demonstrated with an overall high accuracy (88%; CI, 75% - 94%) on predicting the origin site of CUP. However, some tissues such as stomach and esophagus were not still able to be separated by this classifier. Beyond the classifier based on miRNAs or gene expression, few study focus on predicting the origin site of CUP from other epigenetics level, particularly on DNA methylation (DNAm). Since CpG methylation is central to many biological processes and human diseases, which means they are also highly tissue-specific and sensitive for common tumors, so it would be helpful for the detection and prediction of cancer considering DNAm.

In this study, we developed a novel prediction algorithm for three classifier types that can effectively identify highly tissue-specific biomarkers (miRNA or CpGs) from high-throughput miRNA expression or DNAm profiles to evaluate the potential of these data to accurately identify the origin of CUP. A large and comprehensive data set both from miRNA expression and DNAm profiles was obtained from more than 4500 tumor samples and over 500 normal samples, respectively representing 14 commonly recognized sites of origin in the differential diagnosis for CUP. Another considerably prospective meaning of this algorithm is that it also provides us a new perspective to find the potential interaction between selected miRNAs and DNAm both from a same solid tissue.

Our classifier based on miRNA expression and Illumina 450k DNAm profiles are available through the MiRNA-Methylation based <u>C</u>UP’s <u>O</u>rigin <u>P</u>redictor (MMCOP: http://server.malab.cn/MMCOP/) webserver, which enables the researchers to predict the origin site of their interested tumors.

## 2 Methods

### 2.1 Flowchart

Figure 1 is a flowchart of this study, also including the algorithm flow of those three classifiers. After the data preprocessing, 1-level and 2-level feature selection method were respectively applied on miRNA expression and DNA methylation profiles to identify tissue-specific biomarkers, then a random forest algorithm was combined to construct the classifiers.

**Figure. 1.**
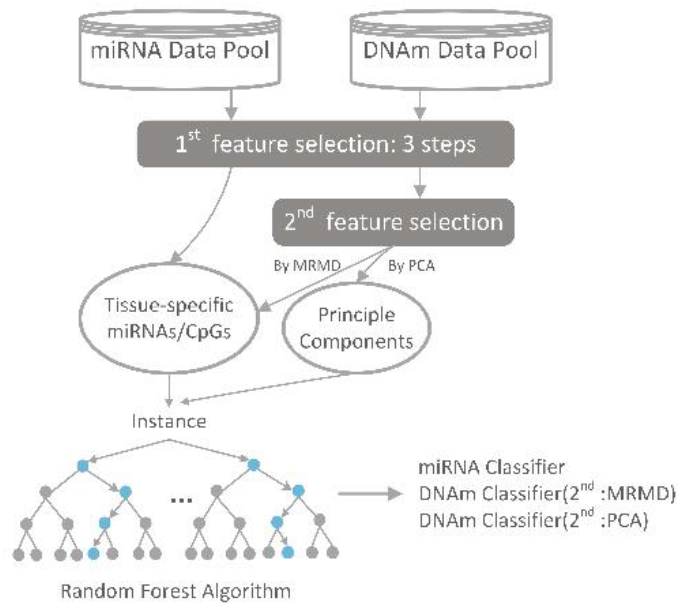
Schematic overview of the workflow of data analysis, and the development of three classifiers.

### 2.2 Tumor samples and normal samples

For data quality control, we made a strict manual review for each data set both in miRNA and DNAm to only select those data meet the requirements of our study. One of the inclusion criterias for subsequent feature selection of miRNAs and DNAm CPGs was that the datasets should have enough samples for both case and control (at least 5) groups. After this manual quality control of dataset, total 6602 samples for miRNA-based profiles, including 6045 tumor samples; total 5379 samples for DNAm-based profiles, including 4668 tumor samples were collected through The Cancer Genome Atlas (TCGA) pilot project (https://tcga-data.nci.nih.gov/tcga/).

The samples based on miRNA expression profiles were sequenced by the BCGSC (IlluminaHiSeq_miRNAseq) sequencing platform, which enable a highly sensitive and specific detection of common miRNAs in human species. And the samples based on DNAm profiles were obtained from Infinium HumanMethylation450 platform, which allowing (for 12 samples in parallel) assessment of the methylation status of more than 480,000 cytosines distributed over the whole genome (Dedeurwaerder, et al., 2011). MiRNA expression and DNAm profiles we selected respectively represented 14 clinically most relevant histologies, covering a broad selection of solid tumors. The details of all tissue samples, including the tumor status, and the histopathologic details of the tumors used for constructing the classifier are provided in Table 1. The details of an independent set of tumors with known origin for validation of the classifier, were also available in this table.

**Table 1.**
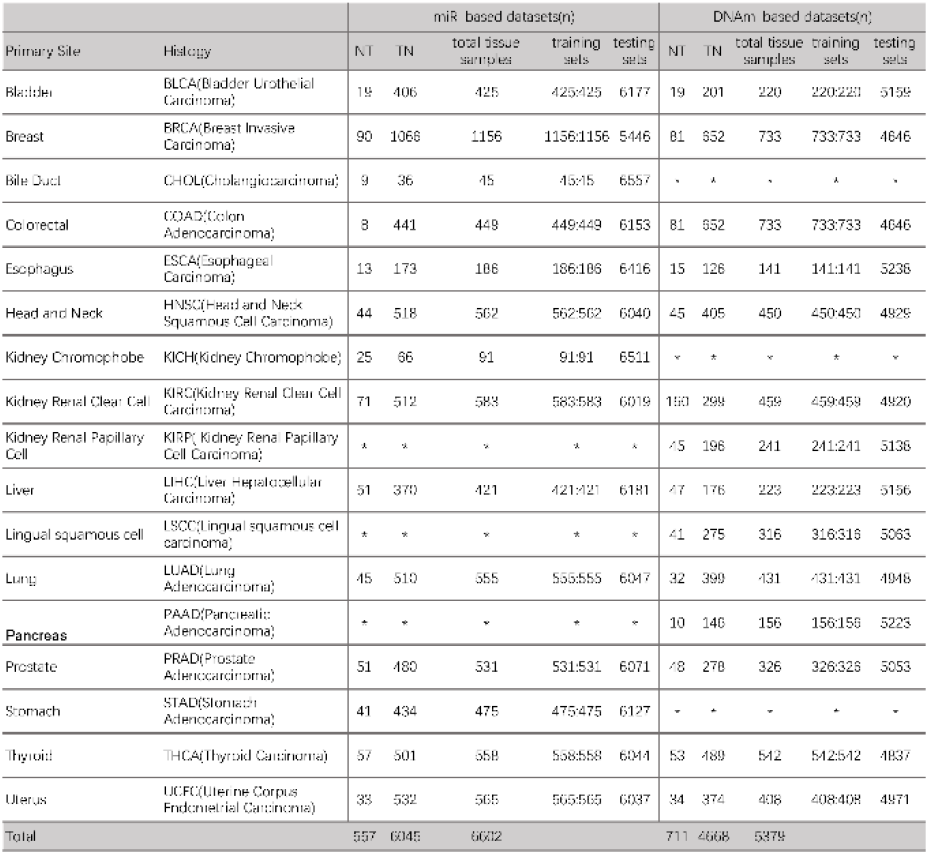
Number of Samples per Tissue for miRNAs expression and DNA methylation profiles, Training sets, Testing sets

**Note**: miR: miRNAs; DNAm: DNA methylation; HeNe: Head and Neck; KRCC: Kidney Renal Clear Cell; KRPC: Kidney Renal Papillary Cell; LSC: Lingual squamous cell; NT: normal tissue sample; TN: tumor tissue sample; Training set: A:A, the 1^st^ A means the total dataset of a given tissue, while the 2^nd^ A means the total datasets of other tissues; *This tissue has not corresponding dataset.

## 2.3. Data Preprocessing and Normalization

All the preprocessing analysis of datasets were performed with the LIMMA package (Linear Models for Microarray and RNA-seq Data, http://www.bioconductor.org/packages/release/bioc/html/limma.html) (Ritchie, et al., 2015) embedded in the R environment (http://www.r-project.org/). For miRNA-based datasets, we selected miRNA isoforms expression data since all isomiRs are from a specific miRNA locus and providing the mature miRNA expression information. For each tissue type of selected datasets, we removed the miRNAs or CpGs which have more than 30% missing value (NA) of the samples. Then, the rest of the missing values are imputed by using the impute.knn function. The maximum miRNA expression value was selected if there were multiple isoforms for a given miRNA in each sample. For DNAm datasets, only one of the multiple samples (technical replication: the sample which was profiled repeatedly by different research organizations) was selected and the absolute methylation values which represent the methylated intensity of every CpGs was calculated by using BMIQ_1.4 (Beta MIxture Quantile dilation) (Teschendorff, et al., 2013) to correct the Type II probe bias. Both for miRNA and DNAm datasets, the expression values of miRNA and the beta value of CpGs were logarithmically transformed (base 2) and quantile normalized.

## 2.4. Feature selection and random forest

### 2.4.1. 1^st^-level Feature Selection

For miRNA-based and DNAm-based samples, 1^st^-level feature selection (three differential analysis steps) was conducted by Limma to identify tissue-specific miRNAs/CpGs and reduce the considerable redundancy of original data. Three different steps were used to select miRNAs/CpGs which should show:

- an abnormally expression/methylated value in a given normal tissue compared with other normal tissue types (one versus all, threshold: *P* − *value* ≤ 0.01);
- the same value in a given cancer tissue when compared with the corresponding normal tissues (one versus all, threshold: *P* − *value* ≥ 0.5);
- an abnormally expression value for the corresponding cancer type when compared with other tumor tissues (one versus all, threshold: *P* − *value* ≤ 0.01).

The 1^st^ step was aimed to select out the miRNAs or CpGs whose mean value level is significantly different between a given normal tissue type and other normal tissue types. Since the goal of this study is to predict the origin of the CUP, the consideration of selecting the biomarkers who have a differential expression level among the different normal tissues would be necessary to help us identify tissue-specific miRNAs or CpGs. Those miRNAs/CpGs with FDR adjusted *P* − *value* ≤ 0.01 were extracted as candidates that have significant differential expression or methylation. For a given tissue dataset which includes both tumor samples and corresponding normal tissue, we also need to assure that the tissue-specific biomarker we select should not have a differentially expressed or methylated value among the different samples (between the normal and cancer samples) which are in the same tissue. So the 2^nd^ step guaranteed that the features we select have no abnormally expression or methylated value in the same tissue. The threshold we set here is 0.5, which means miRNAs/CpGs with FDR adjusted *P* − *value* ≥ 0.5 would be considered a normal miRNA expression or DNA methylation. The 3^rd^ step was used to identify the miRNAs or CpGs who are differentially expressed/methylated among the different tumor tissue types and in order to enable us discriminate cancer types from each other. The threshold we set here was same with the 1^st^ step. And we called this these three steps to select the tissue-specific biomarkers as the 1^st^-level Feature Selection. The interaction of miRNAs and the CpGs which were selected from these three steps would be regarded as preliminary features of 1^st^-level feature selection.

### 2.4.2. 2^nd^-level Feature Selection

For the miRNA-based datasets, the number of selected miRNAs from the 1^st^-level feature selection vary around 10-17. Our subsequent miRNA-based classifier had demonstrated these selected miRNAs are truly tissue-specific biomarkers and are sufficient to enable a high prediction accuracy of CUP. So we don’t need to apply a 2^nd^-level feature selection on the miRNAs selected from the 1^st^-level. However, the DNAm-based datasets sequenced by the platform Infinium HumanMethylation450 have the assessment for the methylation status of more than 480,000 cytosines distributed over the entire genome. Thus, for a certain sample of DNAm profiles, it contains more than 400K CpGs even after the preprocessing procedure. Consequently, due to the extremely large amounts CpGs, the number of the selected CpGs after the 1^st^-level feature selection were still very large and redundant, varying from 10K to more than 20K CpGs for each tissue. Those large feature are still difficult to be used to train the classifier since there must be existing data redundancy. Therefore, we also proposed a 2^nd^-level feature selection to identify much more tissue-specific CpGs. Feature selection method has two main part of the decision: (a) Pearson's correlation coefficient (PCC) is utilized to measure the relevance between features in a subset; (b) Euclidean distance (ED), Cosine distance (CD) and Tanimoto (TO) is utilized to calculate the redundancy among features in a subset. The Pearson correlation coefficient shows the closely relationship between features and labels, while the distance between features is used to present the data redundancy. Thus, we also adopted a java-based method called Maximum-Relevance-Maximum-Distance (MRMD, http://lab.malab.cn/soft/MRMD/index_en.html) (Zou, et al., 2016) which can select features that have strong correlation with labeled and lowest redundancy features subset Pearson. Another commonly used feature selection method is Principal Component Analysis (PCA, embedded in the Dimensionality Reduction part of scikit-learn, http://scikit-learn.org/stable/index.html) (Pedregosa, et al., 2011), which is a statistical procedure, using orthogonal transformation to get the set of values of linearly uncorrelated variables called principal components from observations of possibly correlated variables. Considering the time complexity and whether it is need to do functional annotation of selected CpGs after the feature selection, both these two methods were respectively employed to conduct the 2nd-level feature selection. Both processing of MRMD and PCA have the automatic searching model for the best optimum dimensionality of features. Since the CpGs was selected by MRMD were a set of top optimal number ranked CpGs, which enabled us not only use this feature to construct the classifier, but also try to find out the hided gene information by applying a functional annotation on these CpGs. Whereas, the features selected by the PCA cannot represent the original CpGs since they have been converted to the principle components by an orthogonal transformation, which means we cannot directly adopt a functional analysis for the features of the PCA.

### 2.4.3. Classifier Construction

The miRNA selected from the 1^st^-level feature selection and the CpGs selected from the 2^nd^-level feature selection was used for the tissue-specific biomarkers of every class, which is every tissue as well. All of classes were combined for the further construction of a random forest model. Since selection of relevant biomarkers (e.g., genes, miRNAs, CpG) for sample classification (e.g., to differentiate between patients with and without cancer) is a common research in most genomics studies, another main objective is the identification of small biomarker sets that could be used for diagnostic purposes in clinical practice, which involves obtaining the smallest possible set of biomarkers that can still achieve high prediction performance, in other words, the “redundant” biomarkers should not be selected (Díaz-Uriarte and De Andres, 2006). Therefore, considering the unique characteristics of these research and the properties of genomics data, classification algorithms that be used both for two-class and multiclass problems of more than two classes, or when there are many more variables than observations, and avoid overfit as far as possible would be of great interest for biomarkers classification. Random forest is such an algorithm which has been demonstrated with a high performance on many classification cases of gene microarray (Breiman, 2001; Statnikov, et al., 2008). So here we adopted random forest after the miRNAs and CpGs were selected by the feature selection.

Since the number of the minority class samples (a given tissue class) are very small compared with number of the majority class samples (other tissue classes), it would cause an imbalance problem. To address the imbalanced dataset problem, which may have a serious impact on the performance of classifiers, we adopted a undersampling (Al-Shahib, et al., 2005) method to randomly sample a subset from the majority class to form a balanced dataset with corresponding minority class. Each tissue as well as individual model, which would have a balanced dataset, was trained for the the goal of discrimination of a given tissue from all other tissues (one versus all). For example, in miRNA expression profiles, there are total 425 bladder samples including 19 normal controls and 406 bladder urothelial carcinomas and there are total 6177 other tissue samples, we randomly selected 425 samples from those 6177 samples to construct a balanced dataset via a combination with the 425 bladder samples. The remaining samples from other tissues, as well as the 5752 samples, would be used to test the classifier (see the column 7 or column 12 in Table 1).

Each individual model will be tested by fivefold cross-validation to ensure no over-fitting problem arouse. As a result, there would be a m (m: the number of the types of total tissues) class classifier which included m individual class models. For the classifier testing, we used the remaining datasets via splitting the imbalanced datasets as the testing datasets. So in summary, we proposed a 1-level feature selection + random forest to construct the miRNA-based classifier, while we have 2 options for the DNAm-based classifier: 2-level feature selection (2^nd^ level is MRMD) + random forest or 2-level feature selection (2^nd^ level is PCA) + random forest. A webserver (MMCOP) based on java was also developed to help the users to predict the origin site of tumors by uploading either miRNA expression or DNA methylation profiling.

## 3 Results

### 3.1 Sample Selection

To decide which class of tissues should be included to construct the classifier, we turned to those most metastatic cancers which are most found under a light microscope. The majority CUP, about 90%, are adenocarcinomas, with 60% appearing as moderately to well-differentiated adenocarcinoma, whereas about 30% are poorly differentiated adenocarcinoma. Common adenocarcinoma origins can be found in lung, pancreas, breast, prostate, stomach, liver, and colon tissues (Greco and Hainsworth, 2006; Gupta and Massagué, 2006). About the remaining 10% of CUP can also be found either in squamous cell carcinoma, most of which arise from head and neck tumors, or neoplasms, which are often poorly or even undifferentiated (Greco and Hainsworth, 2006). To ensure a comprehensive representation of the major carcinoma types, defined by their anatomic origin tissue or organ, we selected the major carcinoma (bladder, breast, colon, lung, stomach, kidney, liver, uterus, etc.) and germ cell tumors (clear cell carcinoma). Thus, for miRNA-based and DNAm-based samples, we respectively selected 6602 miRNA samples and 5379 DNAm samples for 14 tissue types which can cover most metastases. Table 1 respectively lists the 14 tissues and histologies (columns 1 and 2).

### 3.2 Feature Selection and Tissue-specific miRNAs expression and CpGs methylation

The key to construct the classifier which has a high performance on predicting origin site of CUP is to use those truly tissue-specific features. So in order to select highly tissue-specific biomarkers, we adopted different strategies for different dataset types. Another consideration of feature selection is the features size for different datasets. For miRNAs-based datasets, which has only about 1800 common miRNAs at first (only 419 miRNAs were left after the data preprocessing procedure), we adopted a 1-level feature which could not only ensure the best quality of identification of tissue-specific miRNAs, but also select appropriate amounts of miRNAs, about 10-17 miRNAs for each tissues with the total of 153 miRNAs selected from the 1^st^ feature selection (see the column 3 in Table 2). However, considering the DNA methylation datasets whose methylation status covers almost all CpGs over the whole genome, which consequently caused the number of CpGs from the 1^st^ feature selection was still large, therefore, we need to adopt a 2^nd^ feature selection to further filter the redundant useless CpGs. Specifically designed for different subsequent processing analysis, we used the MRMD and PCA to respectively process the 2^nd^ feature selection. The optimal number of features selected out for each tissue was determined to by the automatically searching optimum model of MRMD and PCA, since more complex models would include more features without a corresponding increase in subsequent classifier performance. The selected miRNAs with high tissue-specific discriminatory potential from 1^st^ feature selection were listed in the Table 2, also including the number of automatically searching optimal features from both 2-level feature selection. The detailed information of selected CpGs after the 2^nd^ feature selection is available in supplementary materials.

**Table 2.**
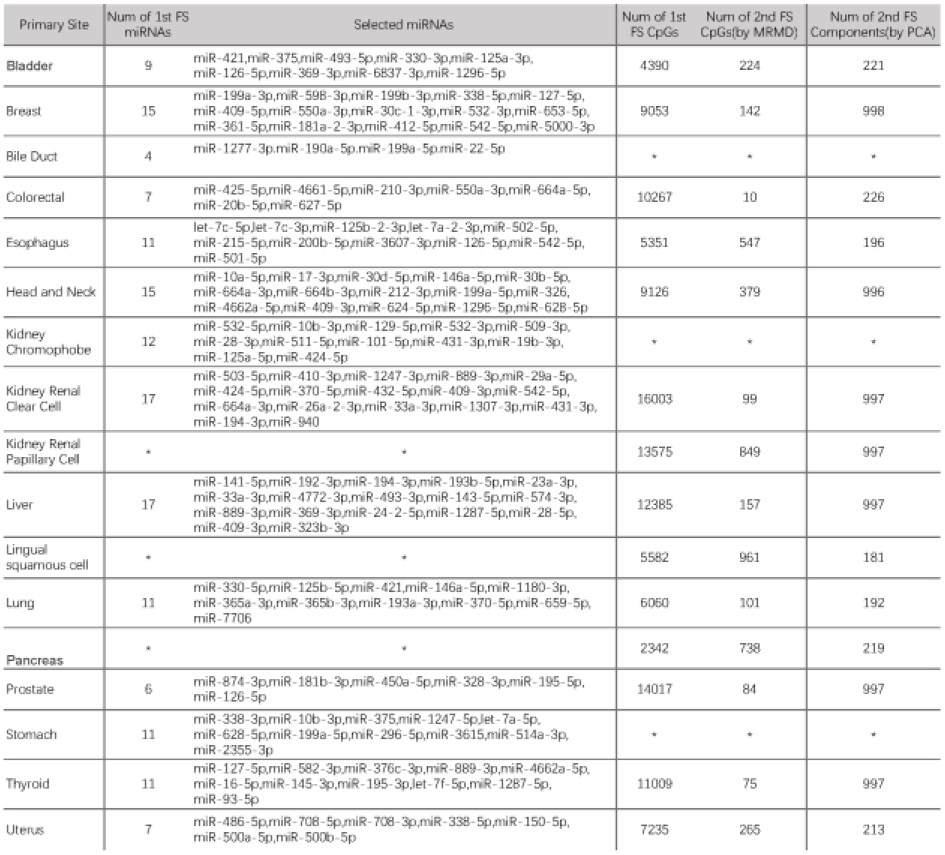
The Number of selected miRNAs and CpGs That Can Be Used for Identification of Tumor Origin, as well as the detailed information of selected miRNAs

**Note**: FS: feature selection; *This tissue has not corresponding dataset; All the miRNAs documented above have omitted the prefix”has-”; The details of CpGs selected by the MRMD are available in the Supplementary Information.

In order to verify the rationality of our feature selection method, we draw a heat-map (Figure 2) of selected tissue-specific miRNAs for the miRNA-based profiles and a box plot (Figure 3) for every top 1 CpGs of 14 DNAm tissues. We can clearly see that some tissues are easy to distinguish from the rest because of a strong and differentially expressed tissue-specific miRNA signature or differentially methylated CpGs.

**Figure. 2.**
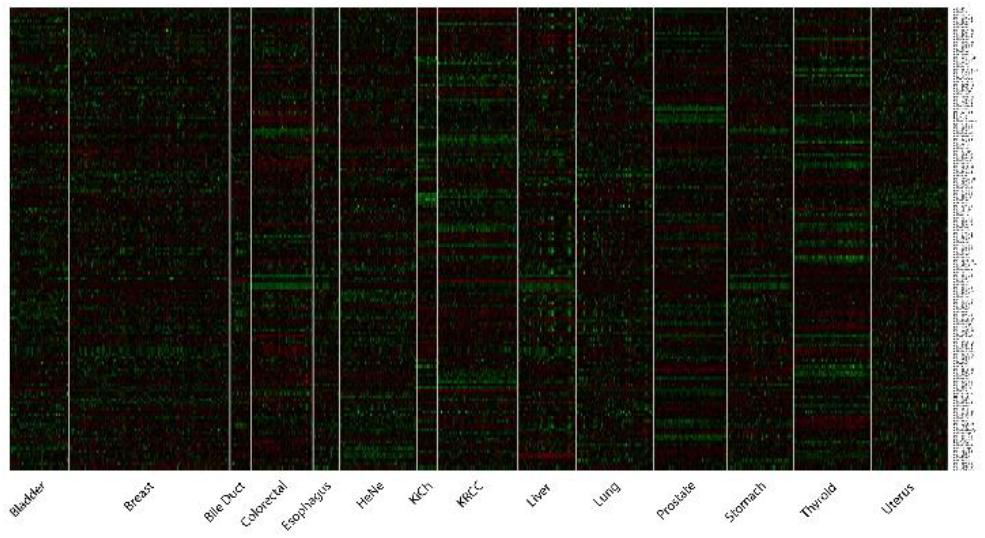
Heatmap of Expression of tumor tissue-specific miRNAs (rows) across 660 samples (columns) that represent the 14 histologies in the training set. These 660 samples were minified in equal proportion from the total 6602 samples. The heatmap shows median normalized log2 data for every miRNA selected in each tissue.

**Figure. 3.**
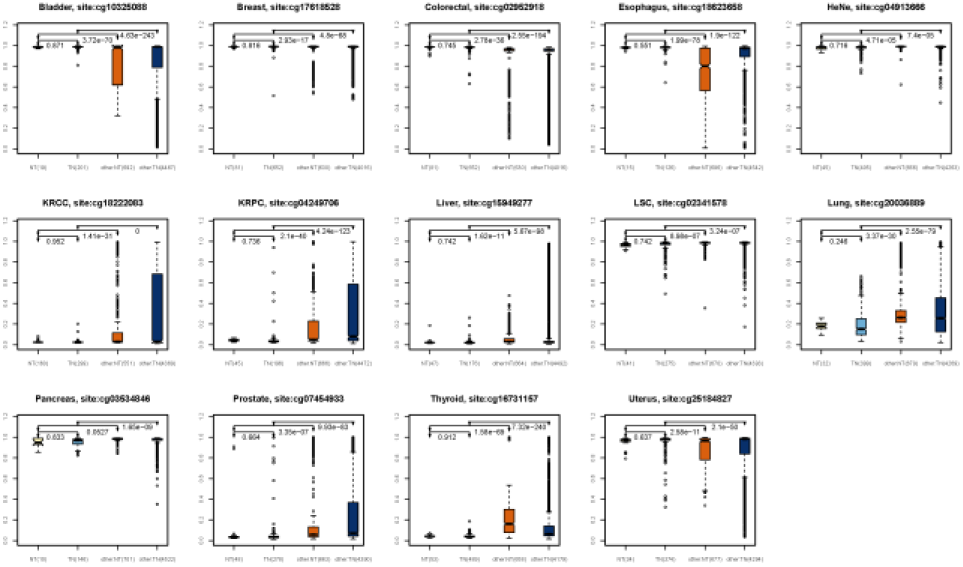
Boxplot of Expression of 14 top 1 tissue-specific CpG site of 14 histologies in the training set. These 14 sites were the top ranking 1st sites after the 2nd-level feature selection; since the boxplot is showing the comparison of a single CpG site, so the p-value between two boxes was calculated by the t-test.

### 3.3 Performance of Classifier on Predicting the Origin of Tumors

In practice, the performance of predictor depends much on the the number of features selected from the feature selection. The optimal number of tissue-specific miRNAs are obtained from the 1^st^ feature selection, while the best performance of DNAm-based classifier obtained using the CpGs selected by the automatically searching optimum model of MRMD and PCA. Specifically, due to the large sample size we selected, we also proposed undersampling method to avoid the potential imbalanced problem. We randomly split the majority class (other tissue samples) into the same sample size as the minority class (the given tissue) to combine a balanced dataset. Then a random forest algorithm would be adopted on this balanced dataset to train the classifier with the optimal selection of tissue-specific biomarkers, an individual model would be consequently generated. The 5-fold cross-validation training accuracy and the testing accuracy on the remaining data (column 7 and column 12 in Table 1) was displayed on Table 3. The tissues which were correctly predicted by either miRNA-based or DNAm-based classifier account for the majority of all cases with respective an overall testing accuracy of 87.78% (miRNA-based, CI, 72.55%-97.54%), 96.43% (DNAm-based by MRMD, CI, 87.85%-99.50%) and 97.06% (DNAm-based by PCA, CI, 92.44%-100%). Specifically, it’s evident that the performance of DNAm-based classifier by PCA is better than the other two classifiers (Table 3).

**Table. 3.**
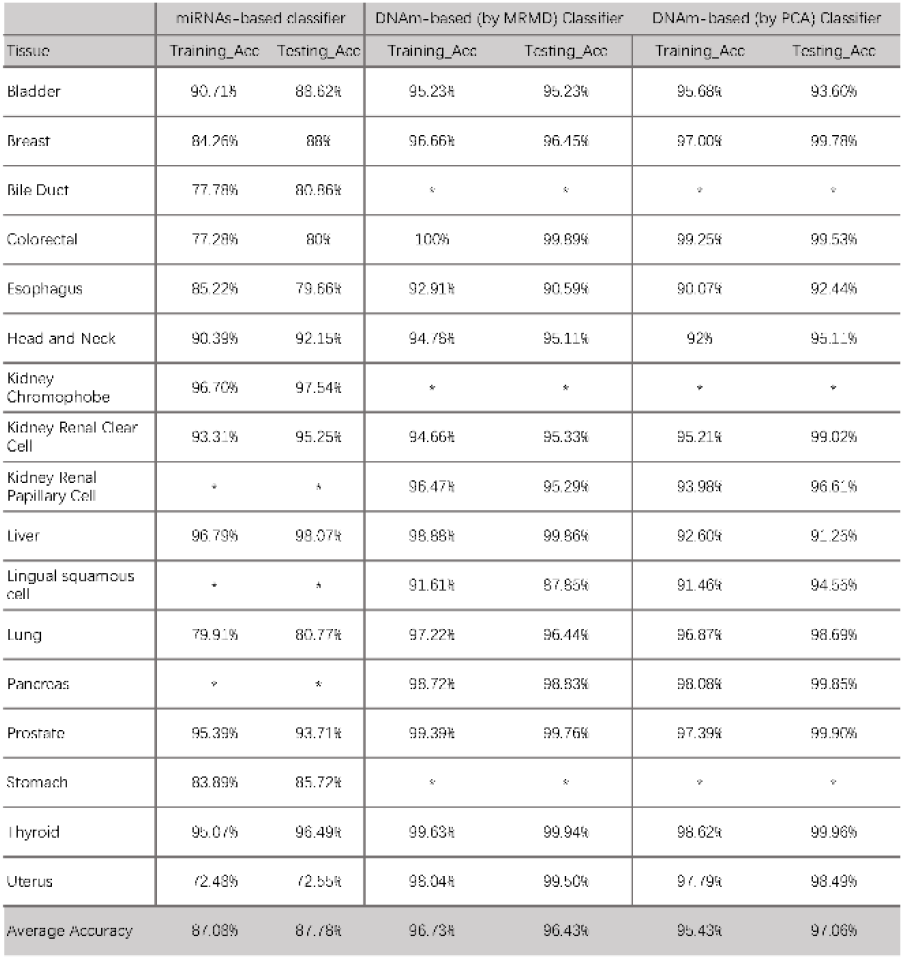
The Performance of Three Classifiers for the miRNA expression and DNA methylation profiles

**Note:** ACC: accuracy; *This tissue has not corresponding dataset.

## 4 Conclusion

Due to the rapid development of NGS, extremely large amounts of sequencing data such as miRNA expression data, methylation sequencing data have been generated and are also becoming more easily available, which also facilitate the development of cancer genomics. Additionally, miRNA expression, as well as CpGs methylation, have been found to show a great association with the origin site of tumor (Hurd and Nelson, 2009; Kim, et al., 2011). Since the efficient treatment of CUP depends critically on the correct identification of the original site, various ways including microscopic techniques (i.e. CT scan), immunohistochemical testing have been proposed to developed to improve the diagnosis of CUP (Kim, et al., 2005). In additional to those techniques, the method based on gene expression profiling which has tissue-specific characteristics may also enable a highly accurate identification of origin of tumor. Though many different algorithms based on machine learning are available for multiclass cancer classification, such as K nearest neighbor, linear discriminant analysis, support vector machine, recursive feature elimination, decision trees, and artificial neural networks have also been proposed from the gene expression profiling (Dudoit, et al., 2002; Guyon, et al., 2002; Khan, et al., 2001; Nutt, et al., 2003; Qu, et al., 2002; Zhu and Hastie, 2004), the characteristic of DNA methylation strong associated with tissue site should also be included to consider constructing the classifier. The beadchip we used here to analyze is the Infinium HumanMethylation450, which is a hybrid of two different chemical assays, the Infinium I and Infinium II assays, called Infinium II, approximately a third of the cytosines are interrogated with Infinium I, but roughly twothirds are interrogated with Infinium II, a chemical assay unavailable in Infinium 27K. Infinium 450K array covers 96% of the CpG islands (CGIs) and also highly presents multiple CGI shores and CpG sites located far from islands (Dedeurwaerder, et al., 2011). Therefore, these methylation profiles, as well as the miRNA expression profiling, especially generated from patients with common disease or cancers, has enabled much potential multitude of information mining which hided in complex human diseases.

In this study, we developed three comprehensive classifier types which include different levels of feature selection direct for different biology data types, including miRNA expression and DNA methylation profiles. Our classifiers for miRNA-based and DNAm-based datasets were both demonstrated with higher prediction accuracy on a spectrum of diagnostically challenging samples compared with other developed classifiers. The miRNA expression and DNA methylation profiles were used to successfully develop three different classification scheme, based on the random forest algorithm, which integrates the consequences of feature selection within the classifier construction.

Mainly in terms of the biology bias, the 1^st^-level feature selection is based on the miRNA differential expression and DNA differential methylation analysis. The 2^nd^-feature selection, either by PCA or MRMD, however, is mainly based on the algorithm from the mathematic methods. Actually, the datasets for detailed analyzed here represents two main data regimes: the first one is that the dimension (D) of data smaller than the samples size (n) (miRNA expression profiles: D=419, n=6602); the second one is the high-dimensional dataset which the ambient dimension of the dimension (D) may be of the same as, or substantially larger than the sample size (n) (DNA methylation profiles: D=395515, n=5379). For the first data regime (D < n), many machine learning-based classifier had been developed to predict the site of tumor origin. Our algorithm which combine the 1^st^-level feature selection (miRNA differential expression analysis) and random forest to construct the miRNA-based classifier has an approximately the same sensitivity as other cancer classification methods. The miRNA-based classifier was evaluated firstly with also a high prediction accuracy 87.08% (CI, 72.48%-96.70%) by five-fold cross-validation on the initial training datasets, and also a high testing accuracy of 87.78% (CI, 72.55%-98.07%). The classifier based on least absolute shrinkage and selection operator (LASSO) algorithm proposed by Rolf et al (Søkilde, et al., 2014) has relative lower prediction accuracy on colorectal class (76.47%, 13 corrected in 17 cases) when it’s compared either with our miRNA-based classifier (80%). Another example is our miRNA-based classifier has an accuracy of 88.62% on predicting bladder tissue, while the K nearest neigh-bor-based miRNA classifier reported by Rosenfeld et al has zero sensitivity to bladder cancer. However, there are still several tissues such as Bile Duct, Esophagus and Uterus, are inherently hard to classify correctly and are often poorly differentiated or dedifferentiated, these three tissues were also reported with a quite low prediction accuracy in other machine-learning methods, such as the method proposed by Rolf et el (Søkilde, et al., 2014). The testing accuracy of these three tissues in our miRNA-based classifier reach respectively about 80.86%, 79.66% and 72.55%, which are not good as other tissues but still better than some other methods, such as the IHC markers method Park et al (Park, et al., 2007) proposed which can be used to identify the cholangiocarcinoma (Bile Duct) wich a quite low accuracy (28%).

The second data regime (D ≥ n) has been much witnessed with the rapid development of data collection technology, which enables more observations to be collected (larger n), and also more variables, as well as the dimensions to be measured (larger D) (Negahban, et al., 2009). Typically, DNA methylation intensity data collected from Infinium 450K array are such data regime. To process this high-dimension data more efficiently, we adopted two widely-used methods (MRMD and PCA) as the 2^nd^-level feature selection to further select the tissue-specific CpGs from the results in the 1^st^-level selection (DNA differential methylation analysis). The overall accuracy has demonstrated that our two DNAm-based classifiers could have a much higher accuracy than miRNA-based classifier (Table 3). The accuracy of five-fold cross-validation on the initial training datasets achieved 96.73% (DNAm-based by MRMD, CI, 91.61%-99.39%) and 95.43% (DNAm-based by PCA, CI, 90.07%-99.25%) and also a very high testing accuracy of 96.43% (DNAm-based by MRMD, CI, 87.85%-99.50%) and 97.06% (DNAm-based by PCA, CI, 92.44%-100%) (see in the Table 3). In addition, almost all of tissues have a very high accuracy more than 90%, also including those tissue with more difficulty to be identified by miRNA expression profiling, such as Uterus, Lung and Esophagus tissues with testing accuracy (98.49%, 98.69% and 92.44%). We can also clearly see that the overall performance of DNAm-based classifier (2^nd^ level is PCA) is better than the DNAm-based classifier (2^nd^ level is MRMD), also much higher than the miRNA-based classifier (except the Liver tissue). However, there’s a drawback for the DNAm-based classifier (by PCA), this classifier could not directly tell us which CpGs were selected out according to the 2^nd^ feature selection.

One of the main challenges in machine learning based classifier development is the effective identification of an appropriate set of features on which a classifier is trained to identify each class accurately. Our two levels feature selection ensure an adequate volume of selected biomarkers to construct an accurate classifier. From Figures 2-3, we could clearly see those biomarkers we selected from the 1^st^ or 2^nd^-level feature selection have strong heterogeneous tissue-specific signatures. Another worthwhile perspective of this study should be highlight is that we actually proposed another novel way to identify those biomarkers selected from the same tissue would have a strong potential association with each other. In other words, we could examine the functional effects of our selected biomarkers through miRNA expression microarray along with methylation profiling. A Gene Set Enrichment Analysis (GSEA) could be applied on the top *m* ranked CpG loci selected from the MRMD method according to the relevance between features and redundancy among features in a subset. It has been reported there exists a strong association between miRNAs and DNA methylation, such as DNA methylation-associated silencing of tumor suppressor miRNAs contributes to the development of human cancer metastasis (Lujambio, et al., 2008; Subramanian, et al., 2005). So one of the major tasks of our future work is that we will try to find out how those potential relationships work between CpGs, miRNAs, targeted genes and the tissue where found them. Through our webserver (MMCOP), users may find the corresponding potential associated miRNAs, CpG loci or targeted gene by submitting their interested biomarkers.

Reverse Transcription Polymerase Chain Reaction (RT-PCR) technique has several advantages than over miRNA expression microarray and DNA methylation array, such as its access to using Formalin-Fixed Paraffin-Embedded (FFPE) tissue (Giulietti, et al., 2001). The ability to accurately predict tumors origin site using a refined number of biomarkers suggests that translation of a miRNA-based or DNAm-based classifier from micro-array to quantitative PCR is possible. Therefore, another our future work will select several biomarkers identified by our method to see whether it would be validated by using existing FFPE tissue.

Taken together, our study suggests that our three classifier types (miRNA-based: DNAm-based by MRMD and DNAm-based by PCA) can efficiently predict the primary origin of tumors, which may help the pathologists to improve the diagnosis and treatment of patients. For most patients with advanced stage CUP, treatments are becoming increasingly specific since treatment trial varies significantly depending on the origin of the tumor. An adjunct genomics diagnostic regimen could enable a more directed clinical evaluation of patients. We believe our classifiers, as well as other classifiers based on relevant biomarkers, such as mRNA and proteins, in combination with clinical investigation will greatly advance and promote rational, specific therapy of patients with CUP through much more focused testing, resulting in reduced patient morbidity, and improved median survivals of patients.

